# Spatial Epigenome Sequencing at Tissue Scale and Cellular Level

**DOI:** 10.1101/2021.03.11.434985

**Authors:** Yanxiang Deng, Di Zhang, Yang Liu, Graham Su, Archibald Enninful, Zhiliang Bai, Rong Fan

## Abstract

Spatial biology is emerging as a new frontier of biomedical research in development and disease, but currently limited to transcriptome and a panel of proteins. Here we present spatial epigenome profiling for three histone modifications (H3K27me3, H3K4me3, H3K27ac) via next-generation sequencing by combining in-tissue CUT&Tag chemistry and microfluidic deterministic barcoding. Spatial chromatin states in mouse embryos or olfactory bulbs revealed tissue type-specific epigenetic regulations, in concordance with ENCODE reference data, but providing spatially resolved genome-wide profiles at tissue scale. Using fluorescence imaging to identify the tissue pixels (20μm) each containing one nucleus allowed us to extract single-cell epigenomes in situ. Spatial chromatin state profiling in tissue may enable unprecedented opportunities to study epigenetic regulation, cell function and fate decision in normal physiology and pathogenesis.

## Main Text

Chromatin state is of great importance in determining the functional output of the genome and is dynamically regulated in a cell type-specific manner (*1-5*). Despite the recent breakthroughs in massively parallel single-cell sequencing(*6-12*) that also enabled profiling epigenomics in individual cells (*13-23*), it is becoming increasingly recognized that spatial information of single cells in the original tissue context is equally essential for the mechanistic understanding of biological processes and disease pathogenesis. However, these associations are missing in current single-cell epigenomics data. Furthermore, tissue dissociation in single-cell technologies may preferentially select certain cell types or perturb cellular states as a result of the dissociation or other environmental stresses (*24, 25*).

Spatially resolved transcriptomics emerged to address this challenge (*26-30*). Recently, we further extended it to the co-mapping of transcriptome and a panel of proteins via deterministic barcoding in tissue (DBiT) (*31, 32*). As of today, it remains unreachable to conduct spatially resolved epigenomics sequencing in a tissue section. Herein, we report on a first-of-its-kind technology for spatial epigenomics named high-spatial-resolution chromatin modification state profiling by sequencing (hsrChST-seq) which combines the concept of in tissue deterministic barcoding with the Cleavage Under Targets and Tagmentation (CUT&Tag) chemistry (*33, 34*) (Fig. 1A and fig. S1). First, a tissue section on a standard aminated glass slide was lightly fixed with formaldehyde. Antibody binds to the target histone modification was added, followed by a secondary antibody binding to enhance the tethering of pA-Tn5 transposome. By adding Mg^++^ to activate the transposome in tissue, adapters containing a ligation linker were inserted to genomic DNA at the histone mark antibody recognition sites. Then, a set of DNA barcode A solutions were introduced via microchannel-guided delivery(*35*) to the tissue section to perform in situ ligation for appending a distinct spatial barcode Ai (i = 1-50). Afterwards, a second set of barcodes Bj (j = 1-50) were flowed on the tissue surface in microchannels perpendicularly to those in the first flow barcoding step. These barcodes were then ligated at the intersections, resulting in a mosaic of tissue pixels, each of which contains a distinct combination of barcodes Ai and Bj (i = 1-50, j = 1-50). The tissue slide being processed could be imaged during each flow or afterward such that the tissue morphology can be correlated with the spatial epigenomics map. After forming a spatially barcoded tissue mosaic, DNA fragments were collected by cross-link reversal and amplified by PCR to complete library construction.

**Fig. 1.**
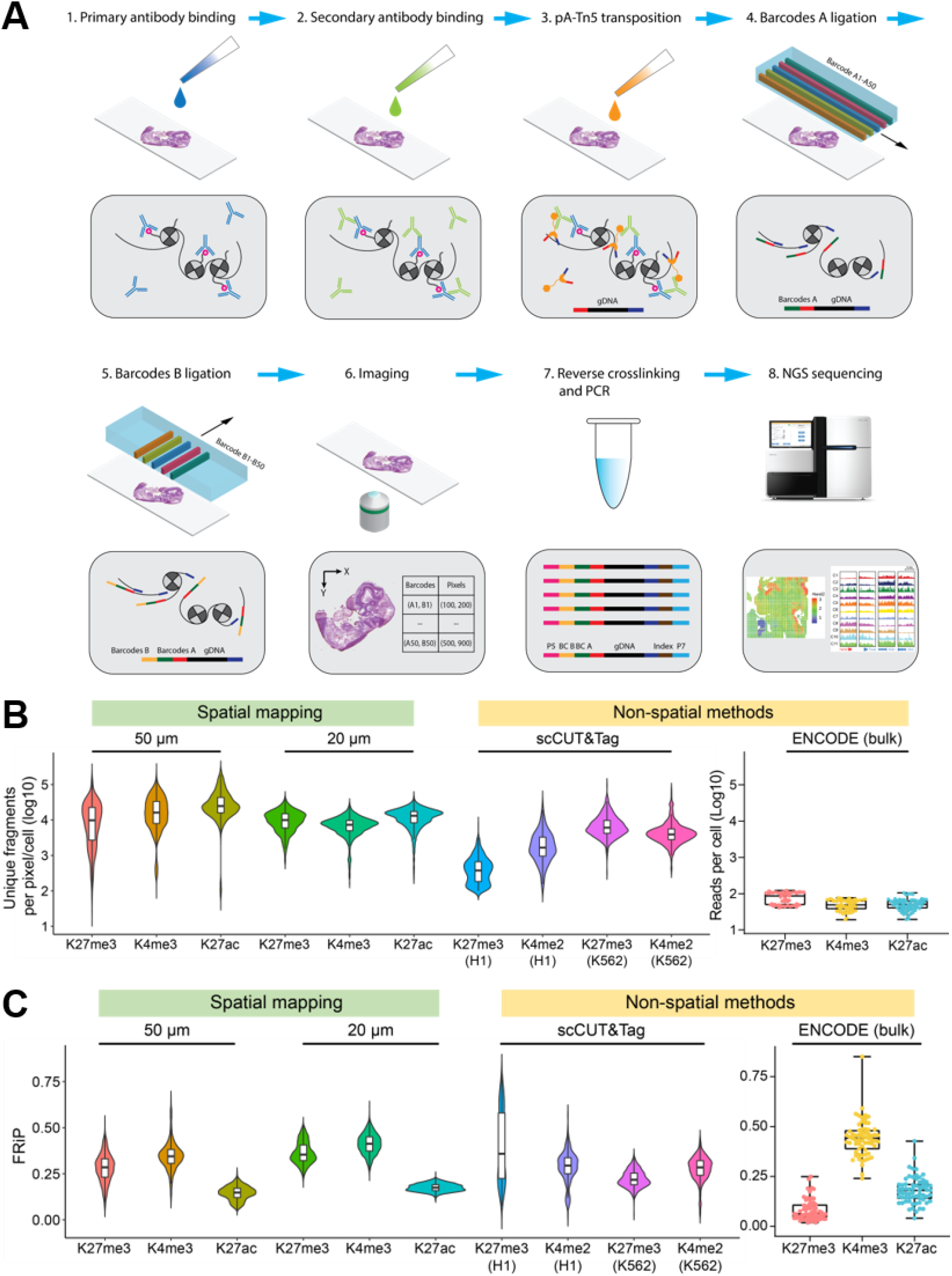
Design of high-spatial-resolution epigenome sequencing of tissue sections. **(A)** Schematic workflow. Primary antibody binding, secondary antibody binding, and pA-Tn5 transposition were performed sequentially in tissue sections. Afterwards, two sets of DNA barcodes (A1-A50, B1-B50) were ligated in-situ. After imaging the tissue sample, DNA fragments were released by reversing cross-linking. Library was constructed during polymerase chain reaction (PCR) and then sequenced by next-generation sequencing (NGS). **(B)** Comparison of number of unique fragments for different histone marks and different microfluidic channel width between our spatial method in this work and other non-spatial chromatin profiling methods. **(C)** Comparison of fraction of reads in peaks (FRiP) for different histone marks and different microfluidic channel width between our spatial method in this work and other non-spatial chromatin profiling methods. Peaks were obtained by peak calling in bulk datasets.

We performed hsrChST-seq with antibodies against H3K27me3 (repressing loci), H3K4me3 (activating promoters) and H3K27ac (activating enhancers and/or promoters) in E11 mouse embryos. We first assessed the quality of spatial epigenome sequencing data based on the total number of unique fragments and fraction of reads in peaks (FRiP) per pixel (Fig. 1B, fig. S3 and fig. S9A). In 50 µm hsrChST-seq experiments, we obtained a median of 9788 (H3K27me3), 16135 (H3K4me3), or 24663 (H3K27ac) unique fragments per pixel of which 28% (H3K27me3), 34% (H3K4me3), or 15% (H3K27ac) of fragments fell within peak regions, indicating high coverage of genomic sequences and a low level of background. In 20 µm hsrChST-seq experiments, we obtained a median of 9951 (H3K27me3), 7310 (H3K4me3), or 13171 (H3K27ac) unique fragments per pixel of which 35% (H3K27me3), 41% (H3K4me3), or 17% (H3K27ac) of fragments fell within peak regions. In addition, the fragment length distribution was consistent with the capture of nucleosomal and subnucleosomal fragments for all modifications (fig. S2, A and B). To validate the reproducibility of hsrChST-seq, we performed correlation analysis between biological replicates. The Pearson correlation coefficient r was ∼0.94 (fig. S2C), which demonstrated a consistent performance of hsrChST-seq. We also compared hsrChST-seq to published non-spatial mapping methods including scCUT&Tag and ENCODE bulk ChIP-seq (*4, 16*), which suggested that hsrChST-seq not only provides spatial information but also better performance and higher data quality.

To identify cell types *de novo* by chromatin states, a cell by tile matrix was generated for the different modifications by aggregating reads in 5 kilobase bins across the genome. Latent sematic indexing (LSI) and uniform manifold approximation and projection (UMAP) were then applied for dimensionality reduction and embedding, followed by Louvain clustering using the ArchR package (*36*). Mapping the clusters back to the spatial location identified spatially distinct patterns that agreed with the tissue histology in a H&E stained adjacent tissue section (Fig. 2, A to C). Cluster 1 (H3K27me3) and cluster 6 (H3K4me3) represent the heart in the mouse embryo. Cluster 2 (H3K27me3 and H3K4me3) and cluster 4 (H3K27ac) are specific to the liver region. Cluster 8 (H3K27me3), cluster 3 (H3K4me3) and cluster 1 (H3K27ac) are associated with the forebrain. Cluster 9 (H3K27me3), cluster 5 (H3K4me3) and cluster 3 (H3K27ac) are the midbrain. Cluster 11 (H3K27me3), cluster 8 (H3K4me3) and cluster 2 (H3K27ac) are the hindbrain. These results demonstrated that hsrChST-seq could resolve major tissue structures with high spatial resolution.

**Fig. 2.**
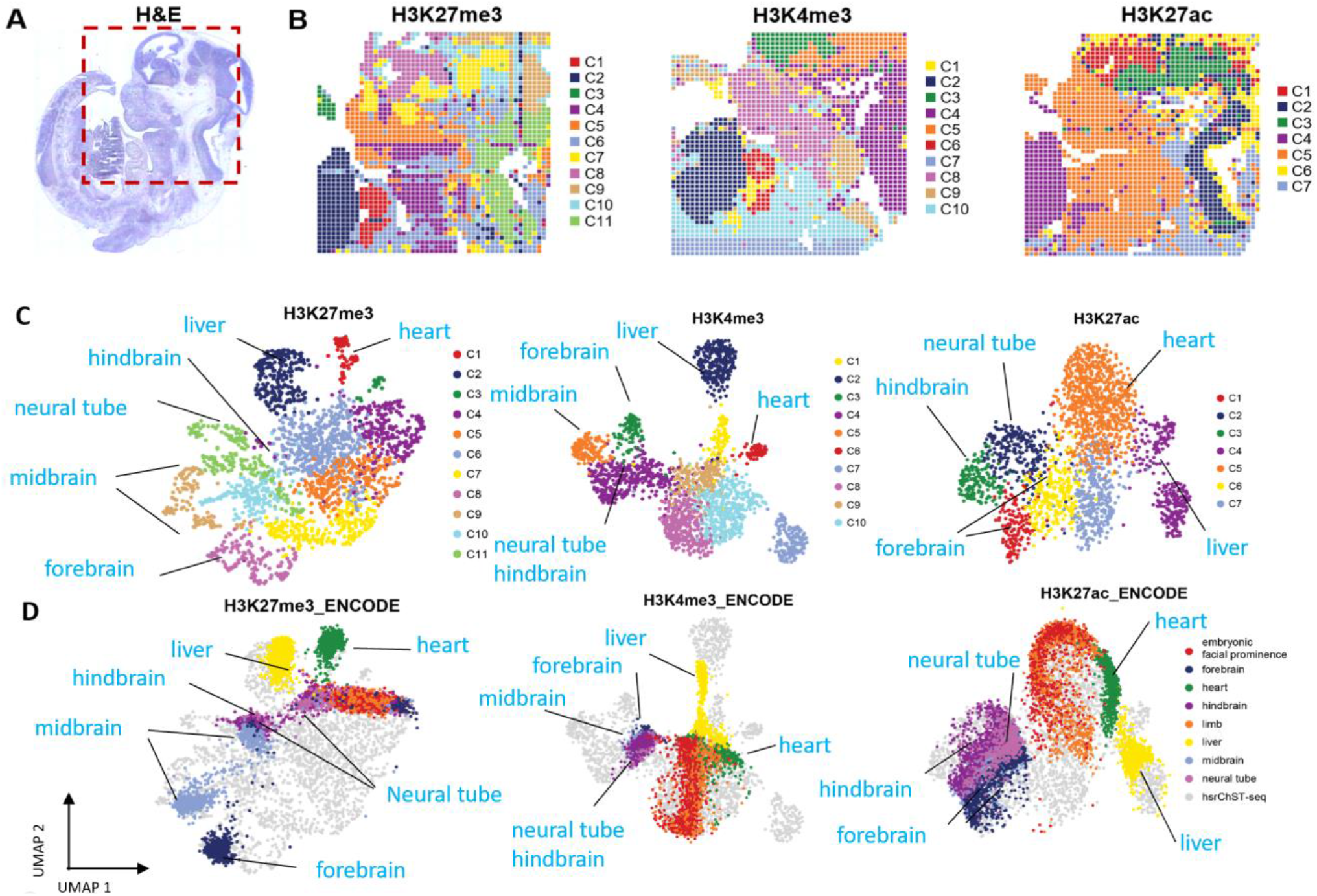
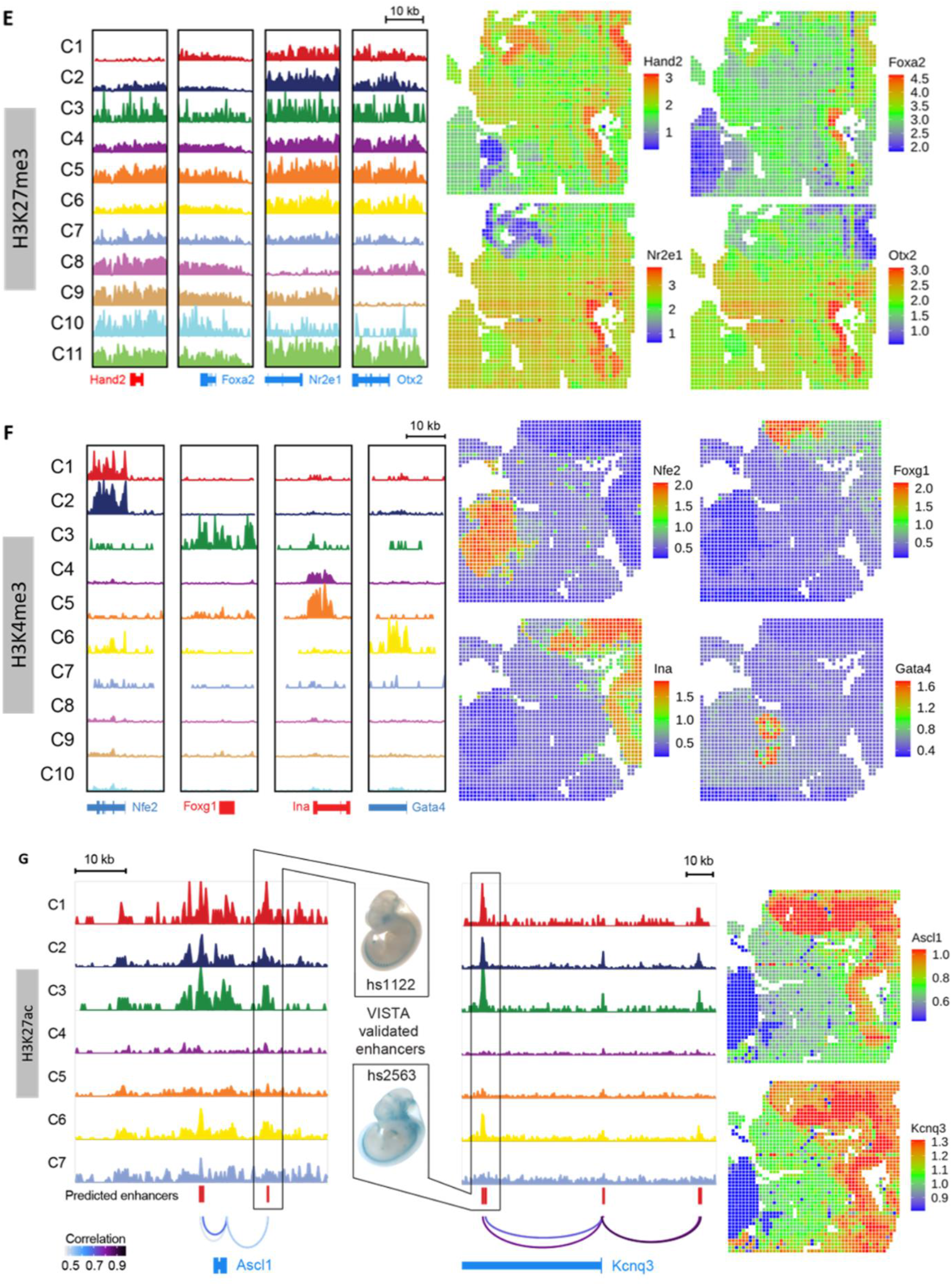
Spatial epigenome mapping of E11 mouse embryos with 50 µm pixel size. **(A)** H&E image from an adjacent tissue section and a region of interest for spatial epigenome mapping. **(B)** Unsupervised clustering analysis and spatial distribution of each cluster for different histone modifications. **(C)** UMAP embedding of unsupervised clustering analysis for each histone modification. Cluster identities and coloring of clusters are consistent with (B). **(D)** LSI projection of ENCODE bulk ChIP-seq data from diverse cell types of the E11.5 mouse embryo dataset onto the hsrChST-seq embedding. **(E)** Genome browser tracks (left) and spatial mapping (right) of gene silencing by H3K27me3 modification for selected marker genes in different clusters. **(F)** Genome browser tracks (left) and spatial mapping (right) of gene activity by H3K4me3 modification for selected marker genes in different clusters. **(G)** Predicted enhancers of Ascl1 (chr10: 87,463,659−87,513,660; mm10) (left) and Kcnq3 (chr15: 66,231,223−66,331,224; mm10) (right) from H3K27ac profiling. Cluster of each track corresponds to (B). Enhancers validated by in vivo reporter assays are shown between main panels.

To benchmark hsrChST-seq data, we used the UMAP transform function to project the ENCODE organ-specific ChIP-seq data onto our UMAP embedding (*4, 36*). Overall, cluster identification matched well with the ChIP-seq projection (Fig. 2, C and D) and distinguished major cell types in E11 mouse embryo. To further compare hsrChST-seq to known spatial patterning during development, we examined cell type-specific marker genes and estimated the expression of these genes from our chromatin modification data. For H3K27me3, chromatin silencing score (CSS) was calculated to predict the gene expression based on the overall signal associated with a given locus (*16*). Active genes should have a low CSS due to the lack of H3K27me3 repressive mark in the vicinity of the marker gene regions (Fig. 2E and fig. S4A). For example, *Hand2*, which is required for vascular development and plays an essential role in cardiac morphogenesis, showed a lack of H3K27me3 enrichment in the heart. *Foxa2*, a transcription activator for several liver-specific genes, had low CSS predominately in the liver region. *Nr2e1*, which correlates with the lack of H3K27me3 modification in the forebrain, is required for anterior brain differentiation and patterning and is also involved in retinal development. *Otx2*, a transcription factor probably involved in the development of the brain and the sense organs, must be highly expressed in the midbrain and hindbrain. For H3K4me3 and H3K27ac, gene activity score (GAS) was used since they are related to active genes (Fig. 2F, fig. S5A and fig. S7A). For example, *Nfe2* and *Hemgn*, which are essential for regulating erythroid and hematopoietic cell maturation and differentiation, were active exclusively in liver. *Foxg1* was highly enriched in the forebrain, which plays an important role in the establishment of the regional subdivision of a developing brain and in the development of telencephalon. *Ina* is involved in the morphogenesis of neurons, showed high GAS in midbrain and hindbrain. *Gata4*, which plays a key role in myocardial differentiation and function, was activated extensively in the heart. We further conducted Gene Ontology (GO) enrichment analysis for each cluster, and the GO pathways matched well with the anatomical annotation (fig. S4B, fig. S5B, and fig. S7B). To understand which regulatory factors are most active across clusters, we calculated transcription factor (TF) motif enrichments in H3K4me3 and H3K27ac modification loci (fig. S6 and fig. S8). As expected, the most enriched motifs in liver correspond to GATA transcription factors, including the well-studied role of *Gata2* in the development and proliferation of hematopoietic cell lineages. *Mef2a*, which mediates cellular functions in cardiac muscle development, was enriched in the heart region. To predict gene regulatory interactions and enhancer target genes across clusters, we correlated scRNA-seq data (*37*) and H3K27ac modifications at candidate enhancers (Fig. 2G). This correlation-based map predicted experimentally validated enhancer-gene interactions with high spatial resolution. For example, predicted enhancers of *Ascl1* and *Kcnq3* were enriched in the brain and eye, which agreed with the VISTA validated elements (*38*).

We then conducted hsrChST-seq with 20 µm pixel size to analyze the brain region of an E11 mouse embryo (Fig. 3, A to C). Unsupervised clustering showed distinct spatial patterns for all modifications, and H3K27me3 identified most clusters. Cluster identification matched ENCODE organ-specific bulk ChIP-seq projection onto the UMAP embedding (fig. S9, B and C). We further surveyed H3K27me3 modifications and observed distinct modification patterns across clusters (Fig. 3, A to C and fig. S10A). *Cfap77* was repressed extensively except in a portion of the forebrain. *Six1*, which is involved in limb development, had low CSS in Cluster 5. Although both *Sfta3-ps* and *Rhcg* lack H3K27me3 enrichment only in the forebrain, they had distinct spatial patterns. Pathway analysis of marker genes revealed that cluster 1 was mainly involved in forebrain development, cluster 2 corresponded to anterior/posterior pattern specification, and cluster 4 was associated with heart morphogenesis, all in good agreement with anatomical annotations (Fig. 3A and fig. S10B).

**Fig. 3.**
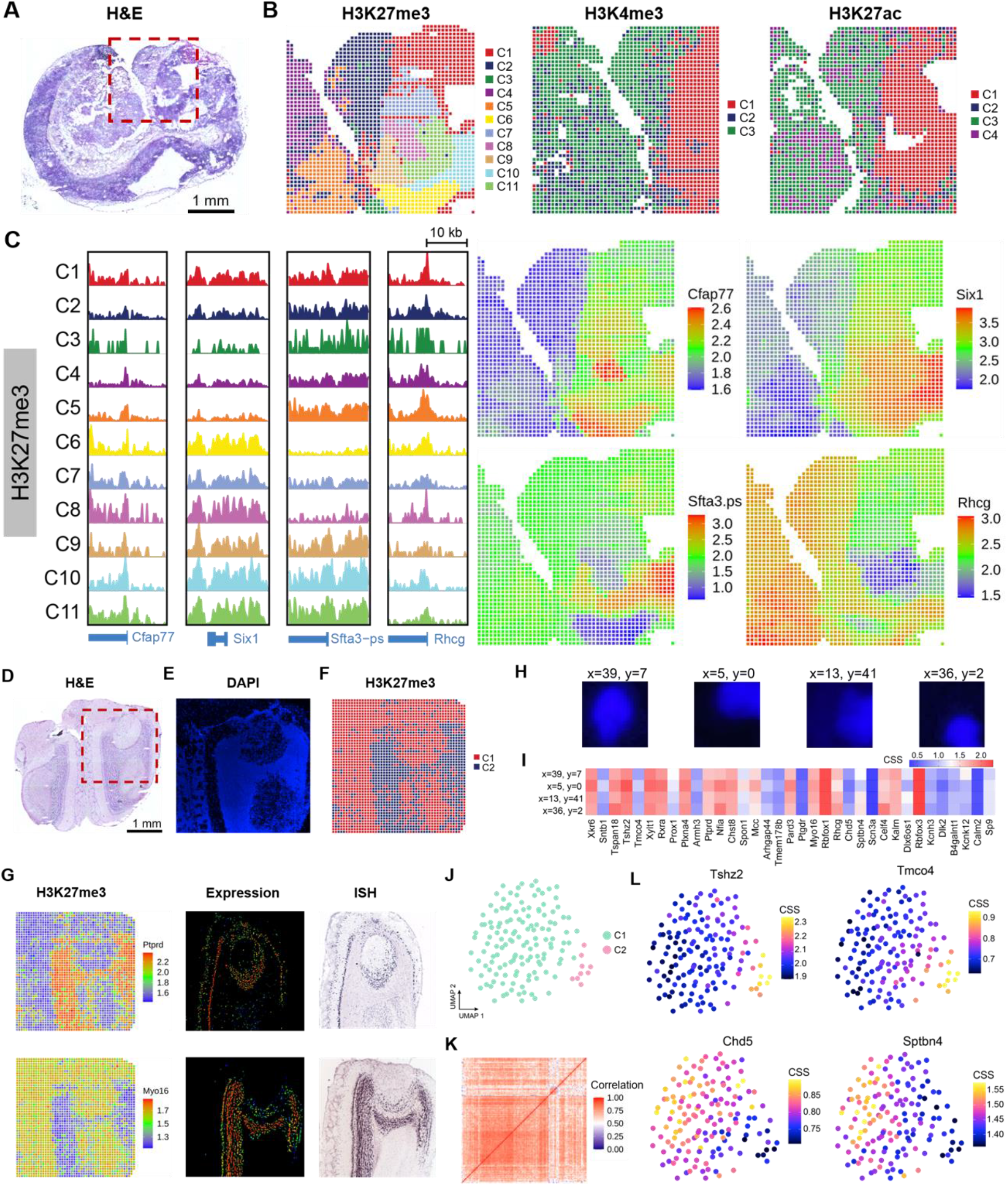
Spatial epigenome mapping at cellular level. **(A)** H&E image of an E11 mouse embryo from an adjacent tissue section and a region of interest for spatial epigenome mapping with 20 µm pixel size. **(B)** Unsupervised clustering analysis and spatial distribution of each cluster of E11 mouse embryo per histone mark. **(C)** Genome browser tracks (left) and spatial mapping (right) of gene silencing by H3K27me3 modification for selected marker genes in different clusters of the E11 mouse embryo data. **(D)** H&E image of mouse olfactory bulb from an adjacent tissue section and a region of interest for spatial epigenome mapping. **(E)** Fluorescent image of nuclear staining with DAPI in a region of interest performed on the same tissue section used for spatial epigenome mapping. **(F)** Unsupervised clustering analysis and spatial distribution of each cluster of mouse olfactory bulb by H3K27me3 modification. **(G)** Spatial mapping (left) of gene silencing by H3K27me3 modification for selected marker genes. In situ hybridization (right) and expression images (middle) of corresponding genes are from the Allen Institute database. **(H)** Fluorescent images of selected pixels containing single nuclei (DAPI). **(I)** Heatmap of chromatin silencing score of selected pixels. **(J)** UMAP of unsupervised clustering analysis of selected pixels containing single nuclei. **(K)** Heatmap of cell-to-cell Pearson’s correlation scores. **(L)** UMAP colored by chromatin silencing score for selected genes.

We also demonstrated hsrChST-seq with immunofluorescence-stained tissue section. A mouse olfactory bulb tissue section was stained with DAPI (4′,6-diamidino-2-phenylindole), a blue nuclear DNA dye (Fig. 3, D and E). Then, we performed hsrChST-seq with H3K27me3, which distinguished the major cell types, including glomerular layer (cluster 1) and granular layer (cluster 2). Examples of H3K27me3 modification patterns revealed by hsrChST-seq and validation by in situ hybridization are shown in Fig. 3G. With DAPI staining for nucleus, we could select the pixels of interest such as those containing only one nucleus or those showing specific chromatin modifications. Combining immunofluorescence with hsrChST-seq at the cellular level (20 µm pixel size) on the same tissue slide allowed for extracting single-cell epigenome data in situ without tissue dissociation (Fig. 3, H to L).

Lastly, to map cell types onto hsrChST-seq data, we integrated the H3K4me3 and H3K27ac data with the scRNA-seq data (*37*). Spatial tissue pixels (black) were found to conform well into the clusters of single cell transcriptomes, enabling the transfer of cell type annotations from single-cell transcriptomics data to the spatial pixels in tissue and further to chromatin modification states. Several organ-specific cell types were detected (Fig. 4 and fig. S11, A and B). For example, the definitive erythroid cells, crucial for early embryonic erythroid development were exclusively enriched in the liver. Cardiac muscle cell types were observed only in the heart region in agreement with the anatomical annotation. Chondrocytes &osteoblasts were observed widely in the embryonic facial prominence. Inhibitory interneurons were highly enriched in the midbrain and hindbrain, and abundant oligodendrocyte progenitors were observed in forebrain region. Although H3K4me3 and H3K27ac had fewer clusters than H3K27me3 in the 20 µm experiments, we found that the clusters that appeared to be homogenous could be further deconvoluted into sub-populations (Fig. 4, E to H). We also co-embedded the H3K4me3 and H3K27ac data with the spatial transcriptome DBiT-seq data from E11 mouse embryos (fig. S11, C and D). In brief, integrative analyses using single-cell or spatial transcriptomics data with well annotated cell types can further refine the definition of cell identity and correlate with spatial distribution of chromatin modification states (*39*).

**Fig. 4.**
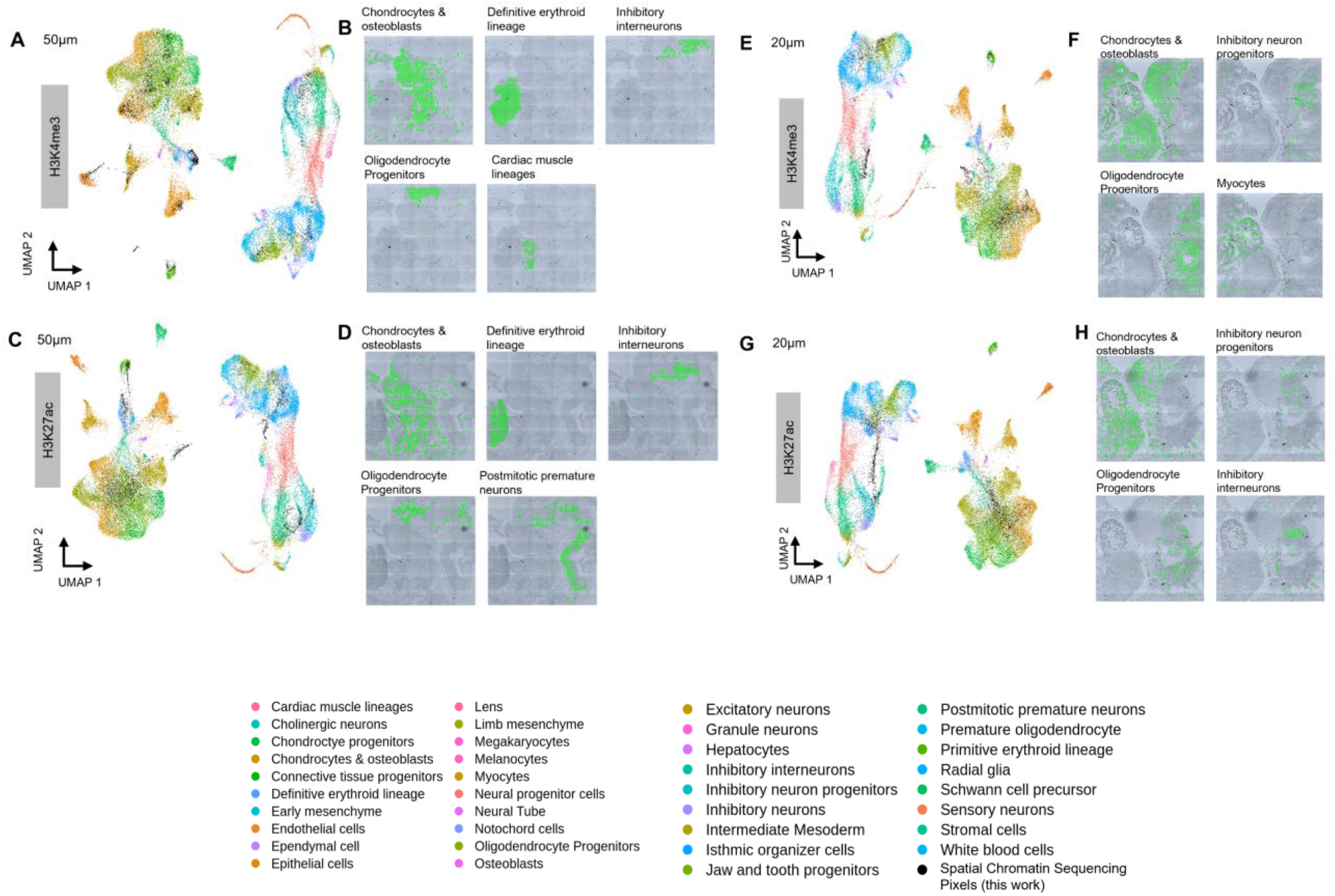
Integrative analysis of scRNA-seq and hsrChST-seq. **(A, C, E and G)** Integration of scRNA-seq from E11.5 mouse embryos (*37*) and hsrChST-seq data. Unsupervised clustering of the combined data was colored by different cell types. **(B, D, F and H)** Spatial mapping of selected cell types identified by label transferring from scRNA-seq to hsrChST-seq. **(I)** List of all identified cell types in UMAP and spatial tissue pixels from hsrChST-seq.

Our study demonstrated the profiling of chromatin states in situ in tissue sections with high spatial resolution. This NGS-based approach is unbiased and genome-wide for mapping biomolecular mechanisms in the tissue context. This capability would enable novel discovery of causative relationships throughout the Central Dogma of molecular biology from epigenome to transcriptome and proteome in individual cells with broad implications for how tissues organize and how diseases develop. The versatility and scalability of this method may accelerate the mapping of chromatin states at tissue scale and cellular level to significantly enrich cell atlases with spatially resolved epigenomics, adding a new dimension to spatial biology.

## Acknowledgments

The molds for microfluidic chips were fabricated at the Yale University School of Engineering and Applied Science (SEAS) Nanofabrication Center. We used the service provided by the Genomics Core of Yale Cooperative Center of Excellence in Hematology (U54DK106857). Next-generation sequencing was conducted at Yale Stem Cell Center Genomics Core Facility which was supported by the Connecticut Regenerative Medicine Research Fund and the Li Ka Shing Foundation.

## Author contributions

Conceptualization: R.F.; Methodology: Y.D., D.Z., and Y.L.; Experimental Investigation: Y.D., D.Z., and Y.L.; Data Analysis: Y.D., and R.F.; Resources: G.S., A.E., and Z.B.; Original Draft: Y.D. and R.F.; Review &Editing: Y.D., G.S., A.E., and R.F.

## Data and materials availability

The sequencing data reported in this paper are deposited in the Gene Expression Omnibus (GEO). Code for sequencing data analysis is available on Github: https://github.com/dyxmvp/hsrChST-seq.

## Supplementary Materials for

**Fig. S1.**
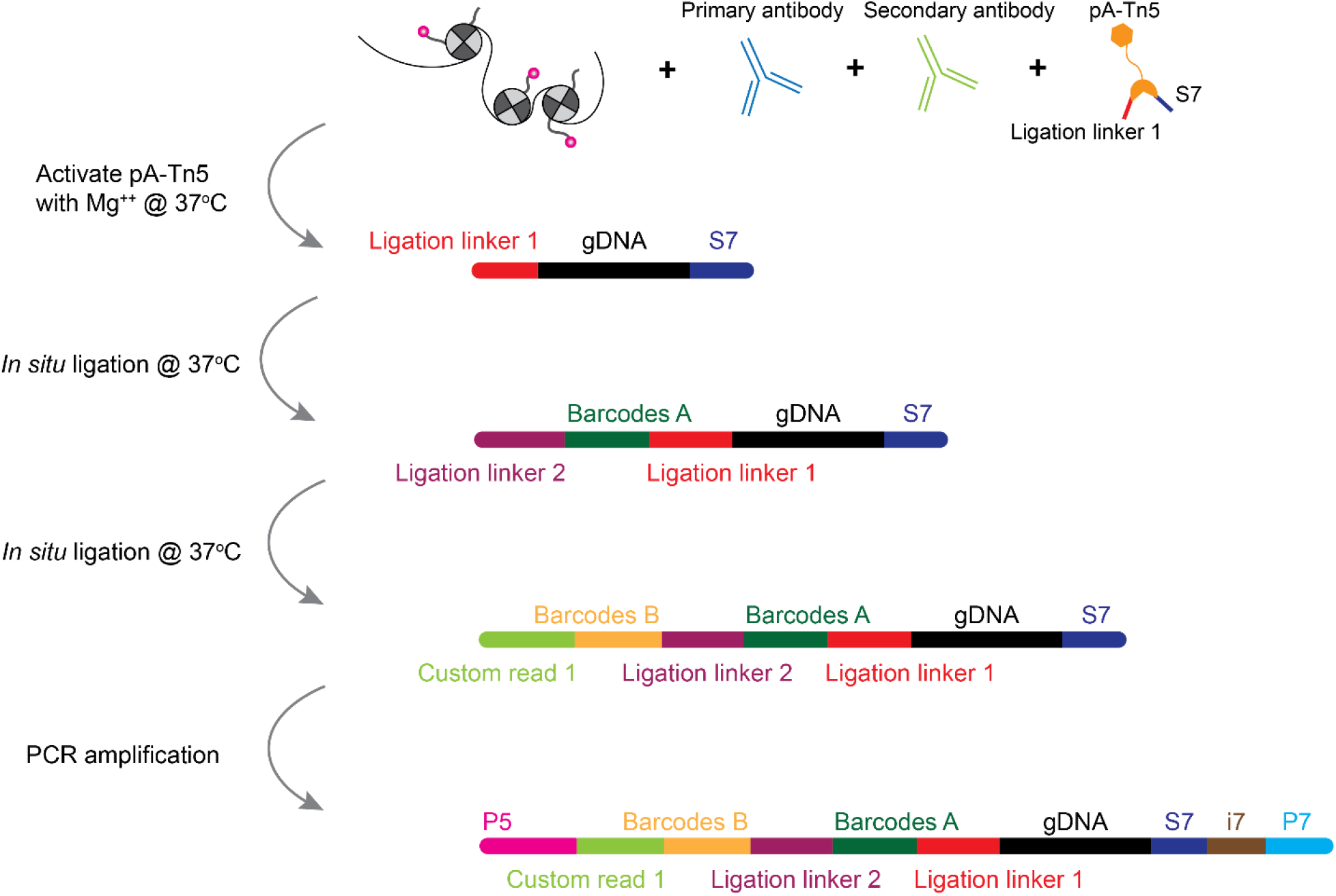
Chemistry workflow of high-spatial-resolution epigenome profiling. A tissue section on a standard aminated glass slide was lightly fixed with formaldehyde. Afterwards, primary antibody that binds to the target histone modifications or chromatin-interacting proteins was added, followed by a secondary antibody binding to enhance the tethering of pA-Tn5 transposome. pA-Tn5 transposome was then activated by adding Mg++ and incubating the sample at 37 °C. Then, the adapters containing ligation linker 1 were inserted to the cleaved genomic DNA at antibody recognition sites. Afterwards, a set of DNA barcode A solutions were introduced by microchannel-guided flow delivery to perform in situ ligation reaction for appending a distinct spatial barcode Ai (i = 1-50) and ligation linker 2. Then, a second set of barcodes Bj (j = 1-50) were introduced using another set of microfluidic channels perpendicularly to those in the first flow barcoding step, which were subsequently ligated at the intersections, resulting in a mosaic of tissue pixels, each containing a distinct combination of barcodes Ai and Bj (i = 1-50, j = 1-50). After DNA fragments were collected by reversing cross-linking, the library construction was completed during PCR amplification.

**Fig. S2.**
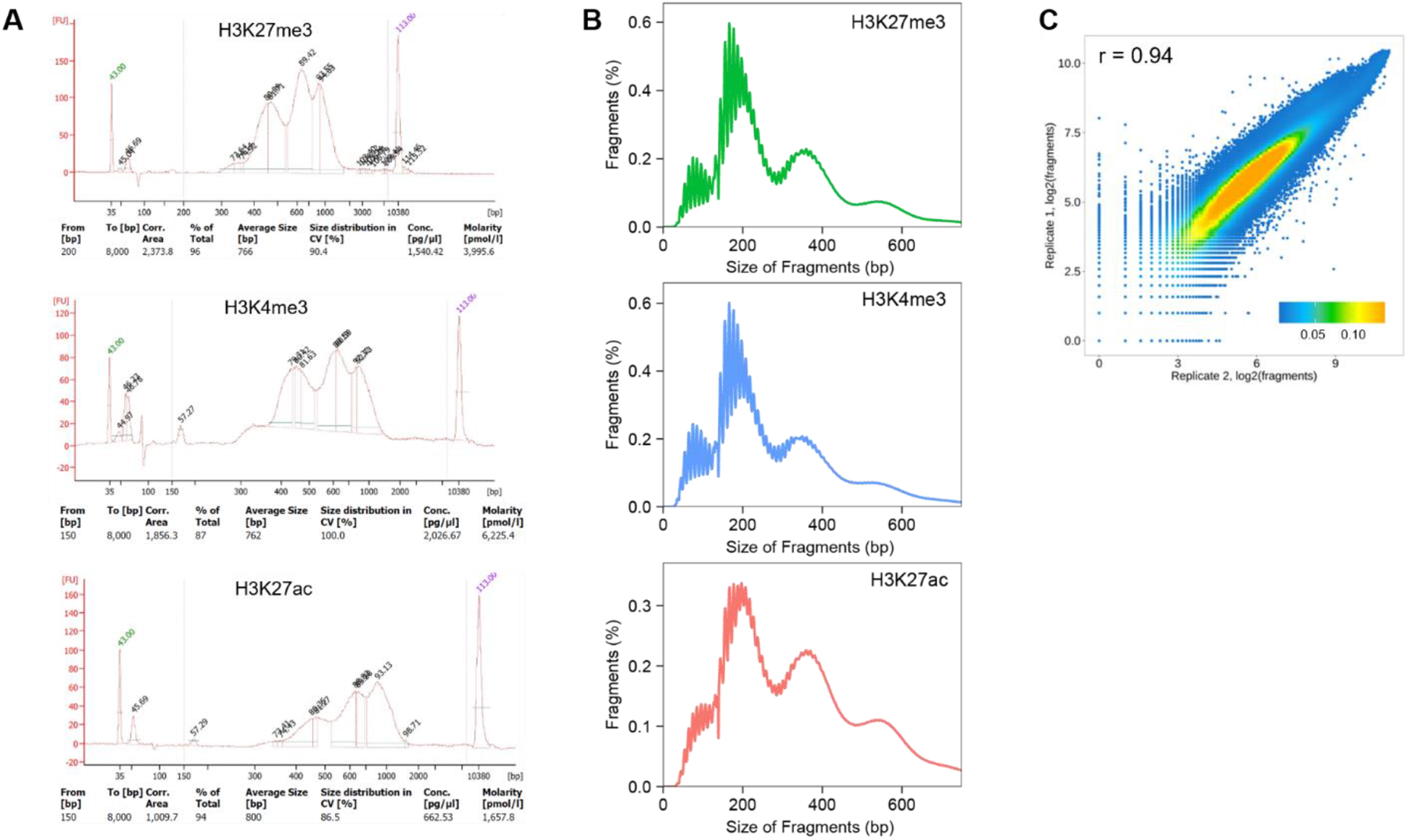
Size distribution of DNA fragments and reproducibility between biological replicates. (A) Bioanalyzer data of DNA fragments. (B) Distribution of fragment lengths. (C) Reproducibility between biological replicates on E11 mouse embryo using H3K27me3 antibody. Pearson correlation coefficient r = 0.94.

**Fig. S3.**
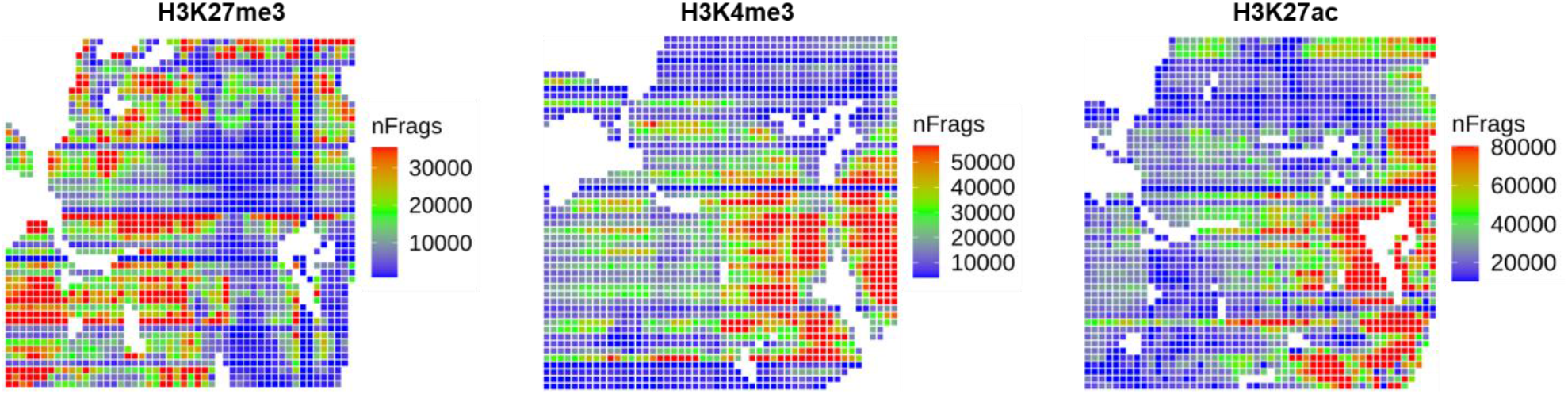
Unique fragment counts in spatial epigenome mapping of E11 mouse embryos using 50 µm devices. These are the spatial heatmaps showing spatial distribution of unique fragment count per pixel analyzed for three different histone marks (H3K27me3, H3K4me3, and H3K27ac).

**Fig. S4.**
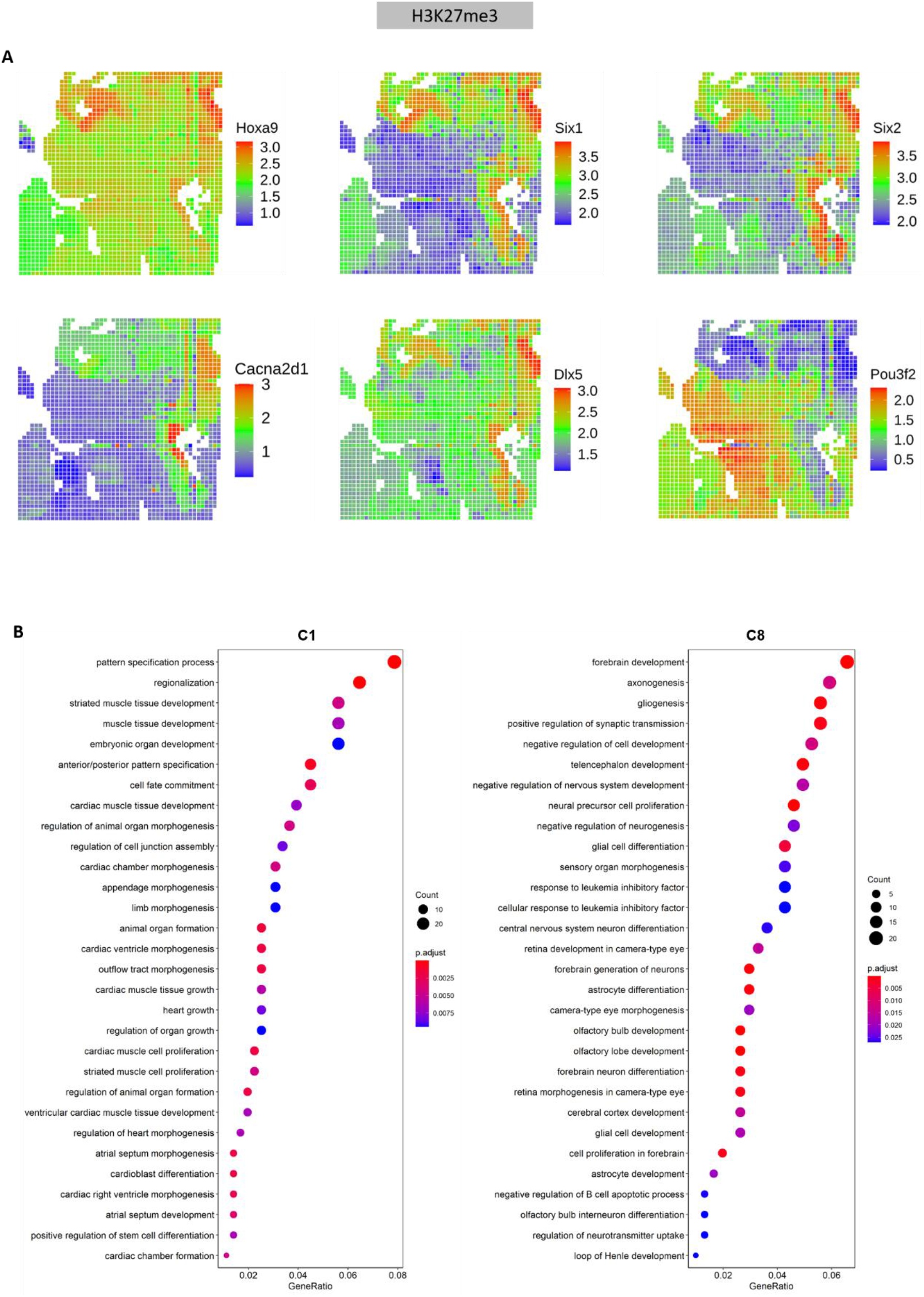
Spatial profiling of H3K27me3 modification of E11 mouse embryos with 50 µm pixel size. **(A)** Spatial mapping of gene silencing by H3K27me3 modification for selected marker genes in different clusters (see Figure 2). **(B)** GO enrichment analysis of differentially silenced genes in selected clusters (C1 and C8).

**Fig. S5.**
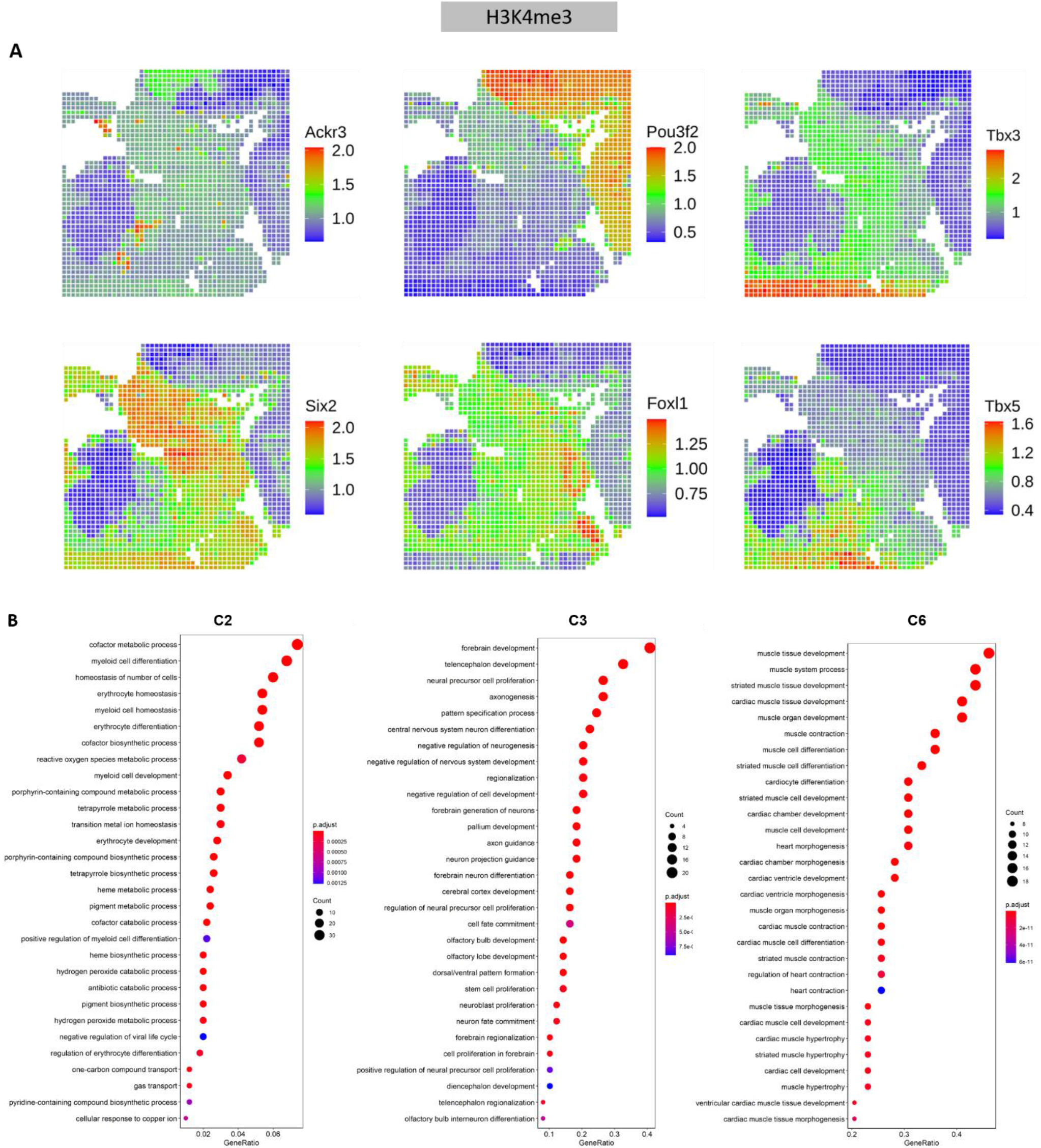
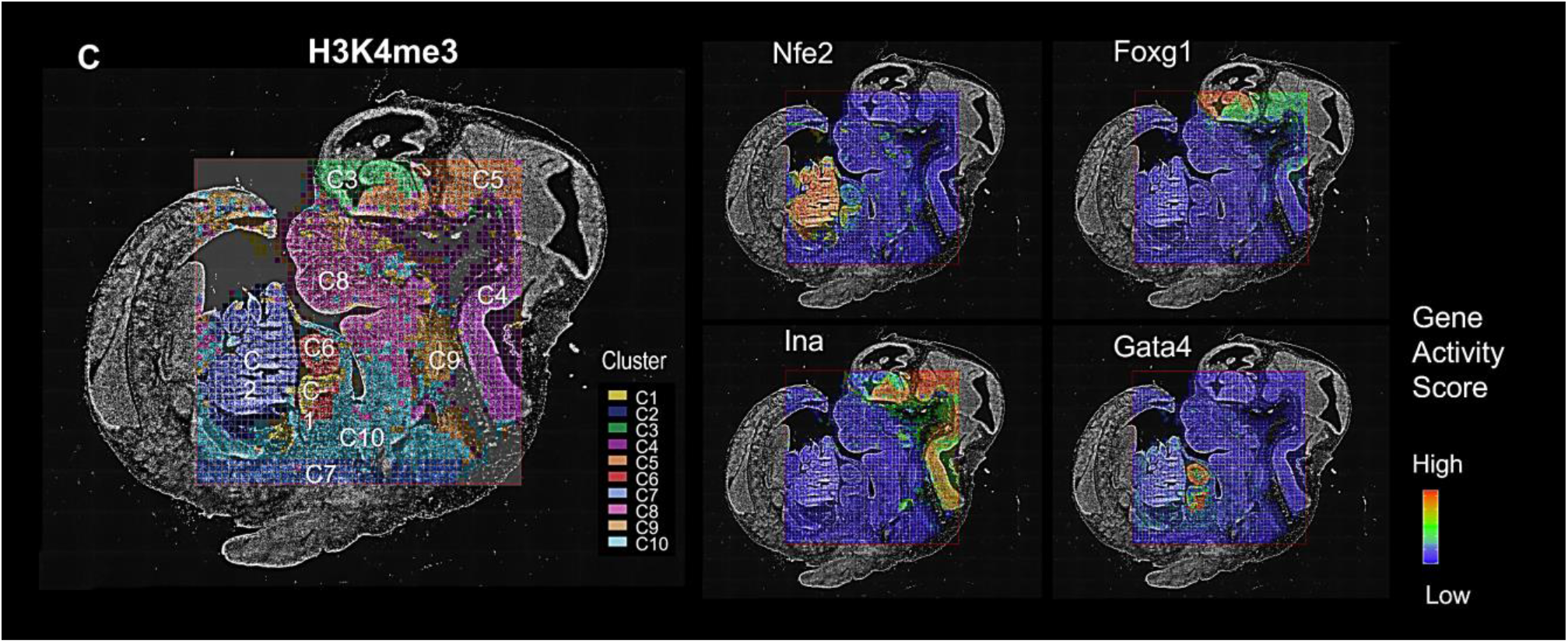
Spatial profiling of H3K4me3 modification in E11 mouse embryos with 50 µm pixel size. **(A)** Spatial mapping of gene activity by H3K4me3 modification for selected marker genes in different clusters (see Figure 2). **(B)** GO enrichment analysis of differentially activated genes in selected clusters (C2, C3, and C6). **(C)** Overlay with the tissue image reveals that the spatial chromatin state clusters precisely match the anatomic regions and the chromatin activity at select genes (*Nfe2, Foxg1, Ina*, and *Gata4*) is highly tissue specific.

**Fig. S6.**
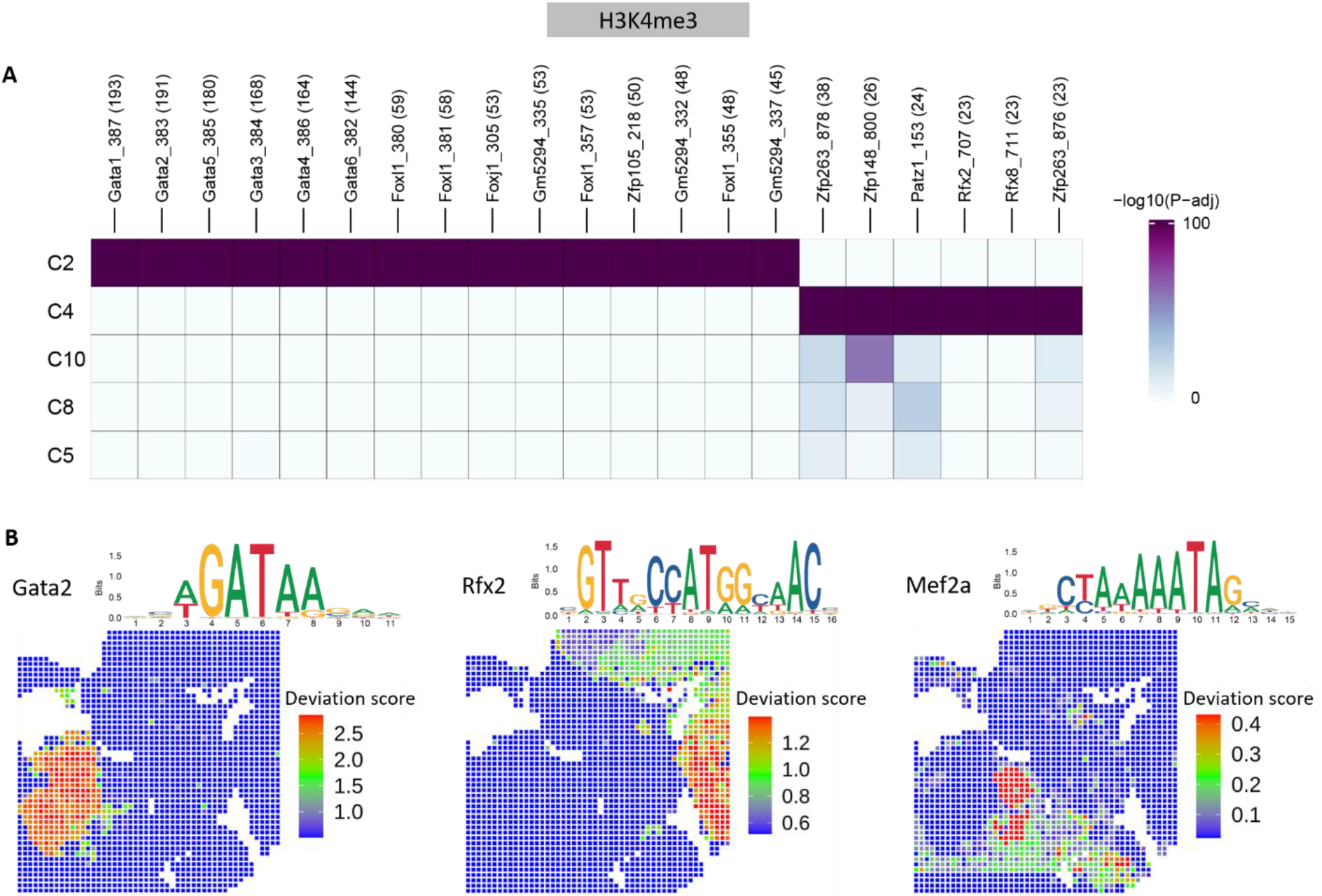
Motif enrichment of H3K4me3 modification in E11 mouse embryos. **(A)** Motif enrichment analysis on marker peaks identified in selected clusters. **(B)** Spatial mapping of transcription factor (TF) motif scores and logo representation of the motif retrieved from the CIS-BP database (*25*).

**Fig. S7.**
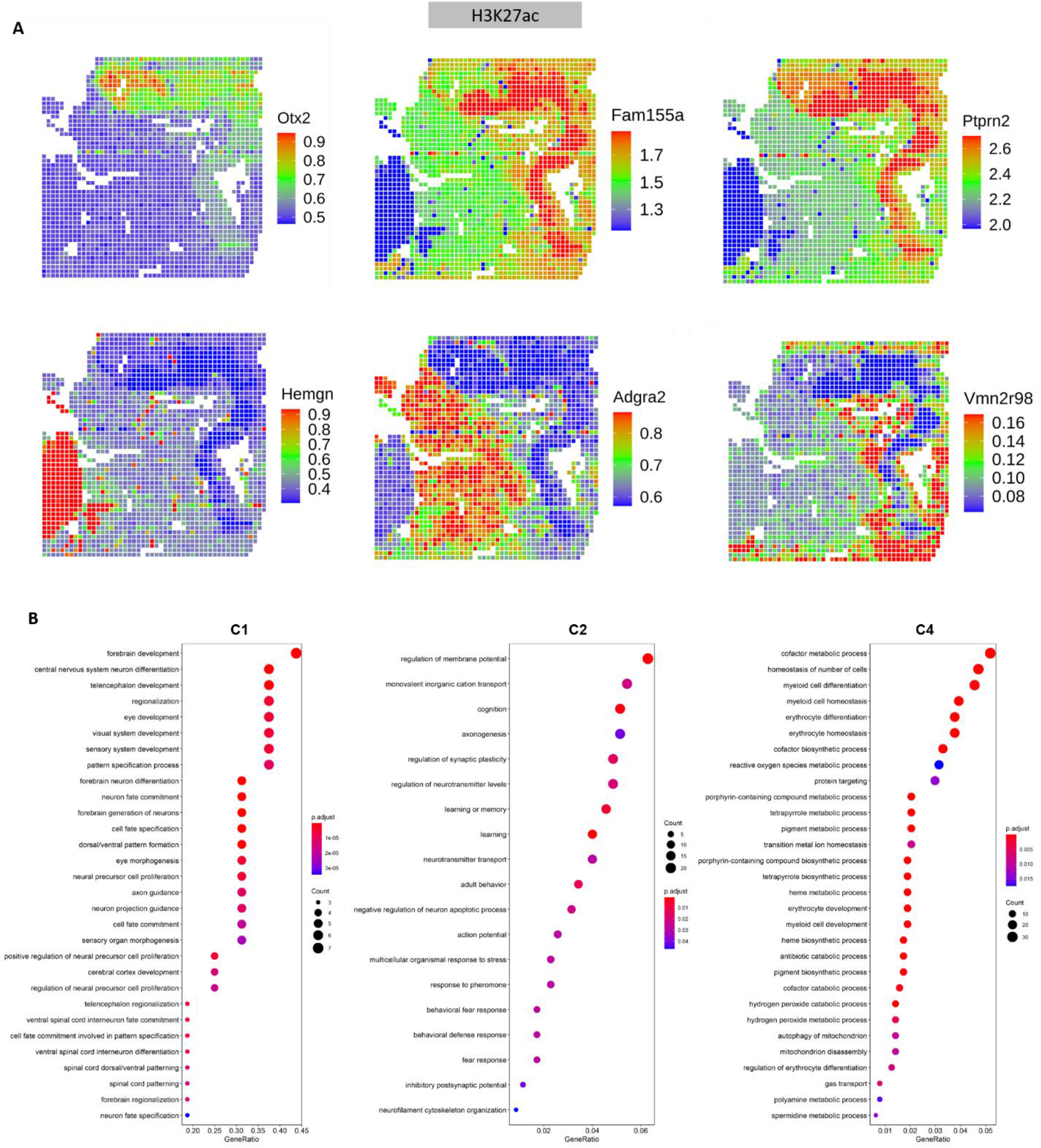
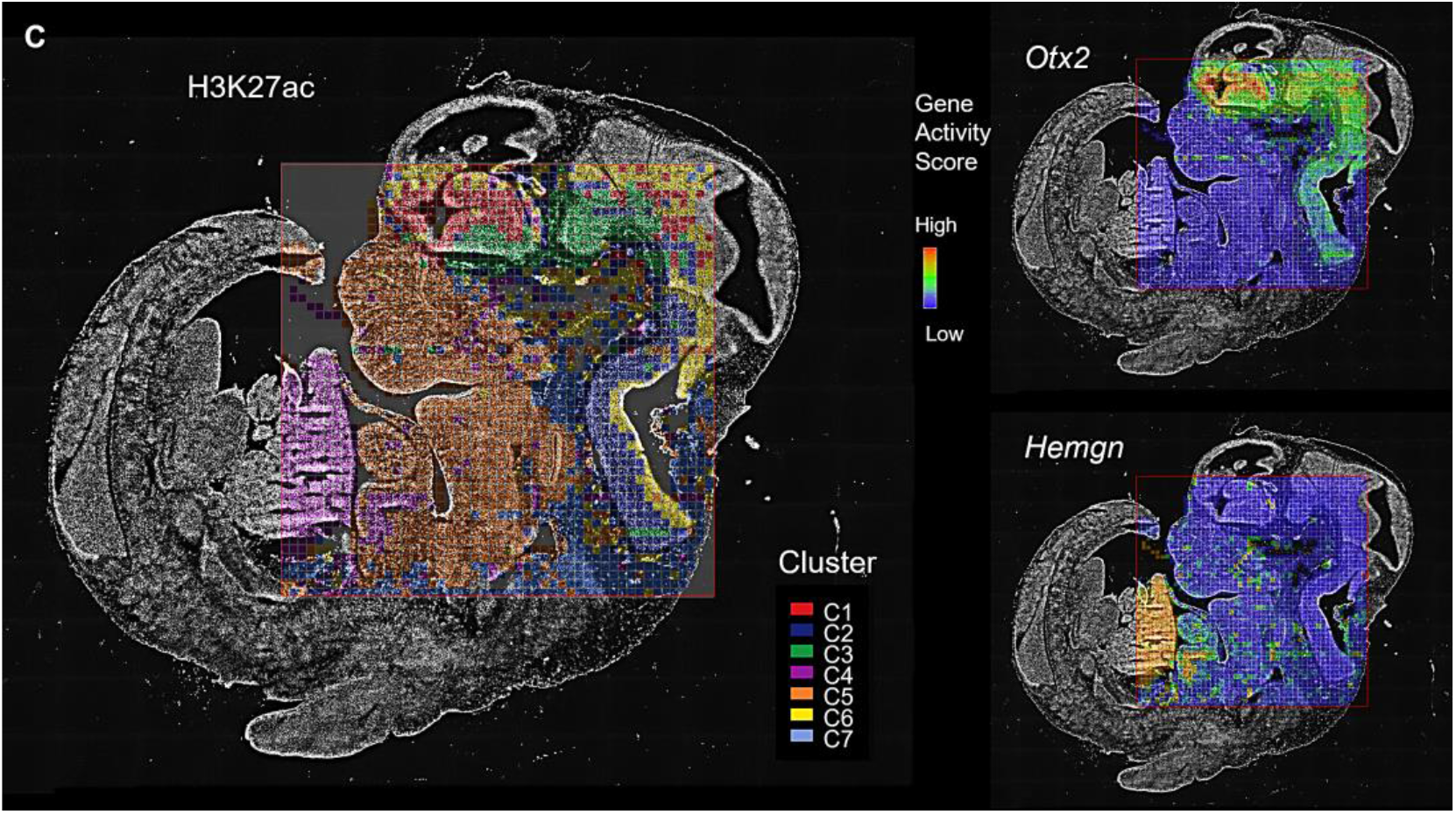
Spatial profiling of H3K27ac modification in E11 mouse embryos with 50 µm pixel size. **(A)** Spatial mapping of gene activity by H3K27ac modification for selected marker genes in different clusters. **(B)** GO enrichment analysis of differentially activated genes in selected clusters (C2, C3, and C4). **(C)** Overlay with the tissue image reveals that the spatial chromatin state clusters precisely match the anatomic regions and the chromatin activity at select genes (*Otx2* and *Hemgn*) is highly tissue specific.

**Fig. S8.**
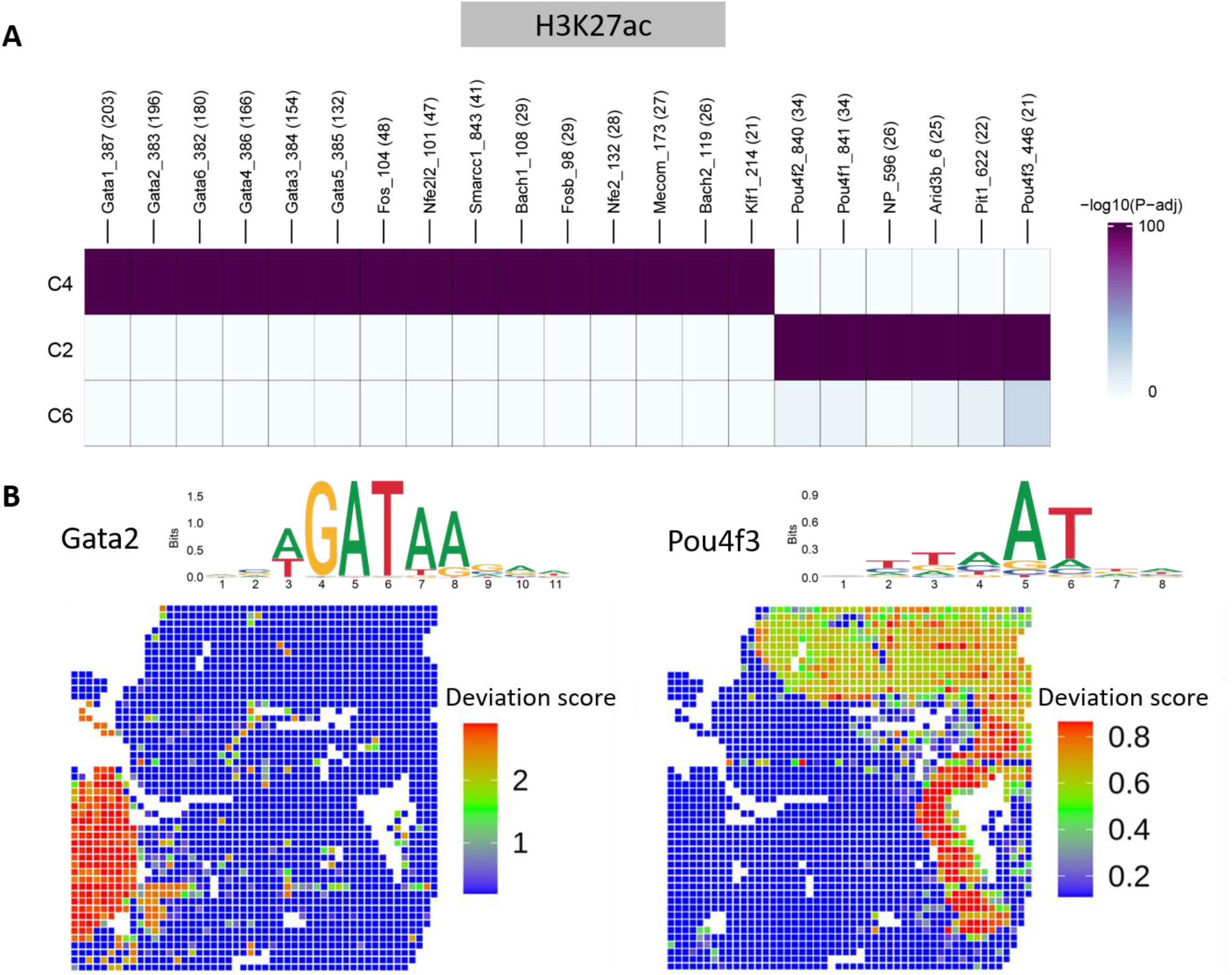
Motif enrichment of H3K27ac modification in E11 mouse embryos. **(A)** Motif enrichment analysis on marker peaks identified in selected clusters. **(B)** Spatial mapping of TF motif scores and logo representation of the motif retrieved from the CIS-BP database (*25*).

**Fig. S9.**
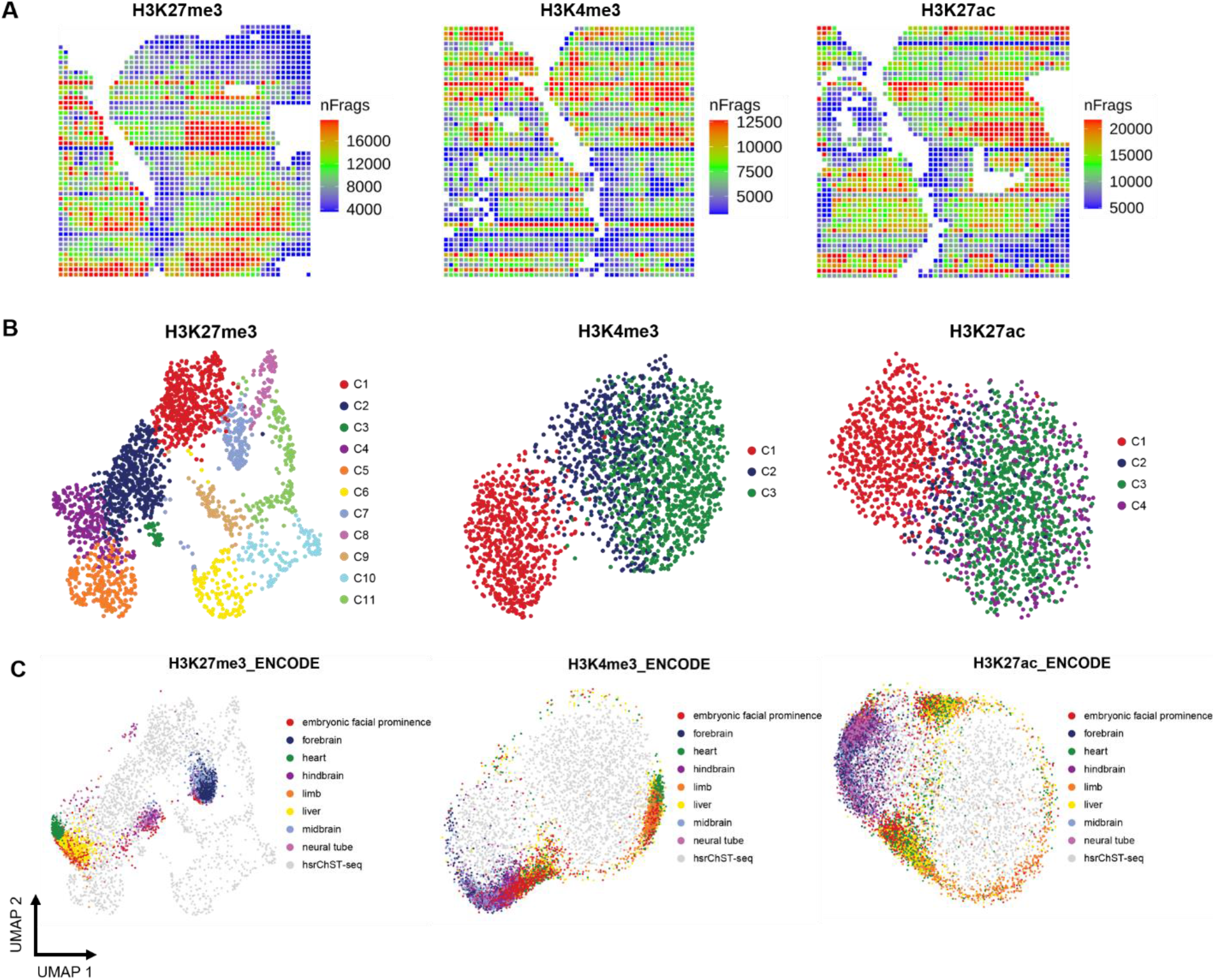
Spatial epigenome mapping of E11 mouse embryos with 20 µm pixel size. **(A)** Spatial maps showing unique fragment count per pixel analyzed for three histone modifications (H3K27me3, H3K4me3, and H3K27ac). **(B)** UMAP embedding of unsupervised clustering analysis for each histone modification. Cluster identities and coloring of clusters are consistent with Fig. 3B. **(C)** LSI projection of ENCODE bulk ChIP-seq data from different organs of the E11.5 mouse embryo dataset into the hsrChST-seq embedding.

**Fig. S10.**
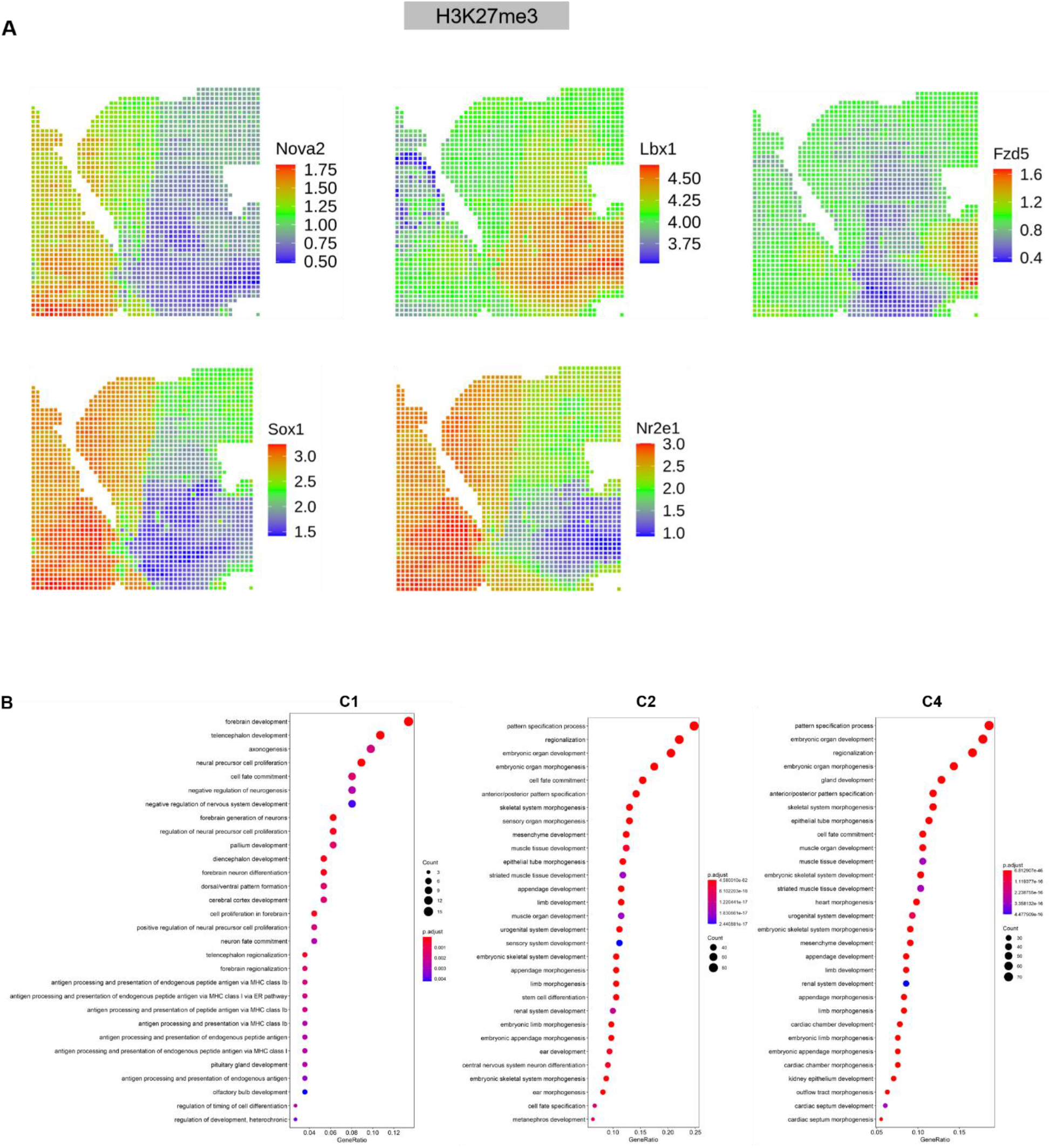
Spatial profiling of H3K27me3 modification in E11 mouse embryos with 20 µm pixel size. **(A)** Spatial mapping of gene silencing by H3K27me3 modification for selected marker genes in different clusters. **(B)** GO enrichment analysis of differentially silenced genes in selected clusters (C1, C2, and C4).

**Fig. S11.**
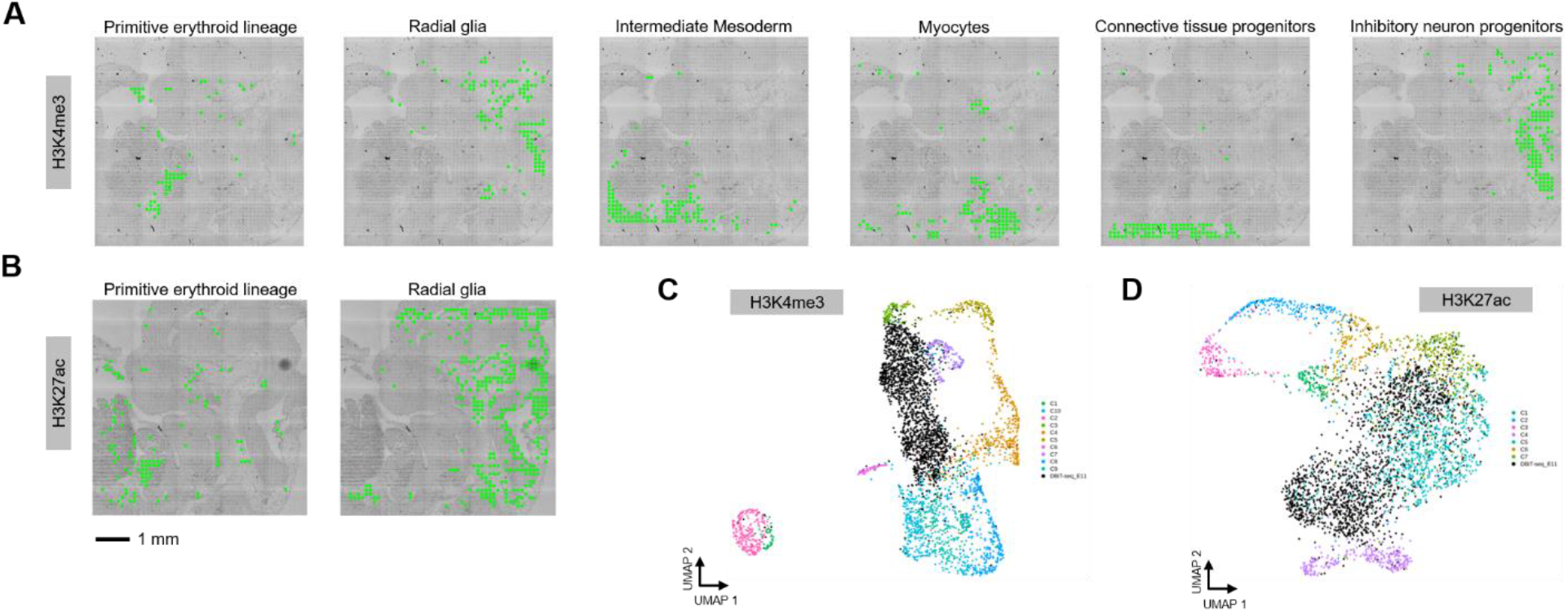
Integrative analysis of scRNA-seq, DBiT-seq and hsrChST-seq. **(A)** Spatial mapping of selected cell types identified by label transferring from scRNA-seq to hsrChST-seq (H3K4me3, 50 µm). **(B)** Spatial mapping of selected cell types identified by label transferring from scRNA-seq to hsrChST-seq (H3K27ac, 50 µm). **(C)** Integration of DBiT-seq from E11 mouse brain (*15*) and hsrChST-seq data (H3K4me3, 50 µm). Cluster identities are consistent with Fig. 2B. **(D)** Integration of DBiT-seq from E11 mouse brain (*15*) and hsrChST-seq data (H3K27ac, 50 µm). Cluster identities are consistent with Fig. 2B.

